# Longevity in plants impacts phylogenetic and population dynamics

**DOI:** 10.1101/2025.06.20.660758

**Authors:** Stephen A. Smith, James B. Pease, Tom Carruthers, Gideon S. Bradburd, Indah B. Huegele, Gregory W. Stull, William N. Weaver, Yingying Yang, Ting-Shuang Yi, Jeremy M. Beaulieu

**Affiliations:** Department of Ecology and Evolutionary Biology, University of Michigan, Ann Arbor, MI 48109, USA; Department of Evolution, Ecology and Organismal Biology, The Ohio State University, Columbus, OH 43210, USA; Idaho Museum of Natural History, Idaho State University, Pocatello, ID 83209, USA; Department of Botany, National Museum of Natural History, Smithsonian Institution, Washington, D.C. 20560, USA; Kunming Institute of Botany, Chinese Academy of Sciences, Kunming, Yunnan 650201, China; Department of Biological Sciences, University of Arkansas, Fayetteville, AR 72701, USA

**Keywords:** simulations, agent-based models, phylogeny, substitution rates, longevity, life history

## Abstract

Many long-lived plant species exhibit notable patterns in phylogenies, such as short molecular branch lengths and high gene-tree conflict. However, it is not clear what biological properties of long-lived plant species or concomitant processes acting within these lineages generate these patterns. To explore this mystery, we implemented an agent-based model and conducted simulations to investigate how longevity affects molecular evolution and population dynamics. Through these simulations, we demonstrated that the patterns exhibited in empirical datasets for long-lived species can be explained by their lifespan and overlapping generations. We also show that somatic mutations can exacerbate these patterns, although evidence for substantive rates in empirical systems high enough to impact phylogenetic patterns is scarce. We discuss several empirical datasets containing life history shifts that exhibit diverse phylogenomic patterns. The variation produced through different parameterizations of our simulations reflects the diversity of patterns found in empirical datasets. Our results have broad implications for phylogenomic patterns and population genetics in general, as well as for specifically explaining patterns of evolution in long-lived lineages.

## Introduction

Life histories across plants range from the ephemeral to the effectively eternal. In natural populations, beech trees (Fagus) set their first seeds after 40 years and can live for hundreds of years; Arabidopsis individuals live their entire life within six to twelve weeks. Variation in life histories can cause markedly different selection pressures leading to dramatically different adaptations (Salguero-Gómez et al. 2016; Chien et al. 2023). For example, a thousand-year-old oak will likely experience the upheaval of several large-scale climate events while concurrently subject to the tides of seasonal variation.

Diverse life history strategies and evolutionary transitions can pose substantial challenges to phylogenetic inference (Smith & Donoghue, 2008; Lanfear et al. 2013). Smith and Donoghue (2008) demonstrated that substitution rates were slower for woody lineages and discussed longevity as a potential cause. Expanding on earlier studies primarily focused on a few “neutrally evolving” gene or spacer regions, Yang et al. (2012) demonstrated that substitution rates were lower for longer-lived species than shorter-lived species across thousands of genes within the Caryophyllales. Lanfear et al. (2013) showed that life-history traits, including generation time, woodiness, and height, are significantly associated with substitution rates across angiosperms. Smith and Beaulieu (2009) further demonstrated associations with molecular rate and climate evolution. Beaulieu et al. (2010) found that rates of genome size evolution also vary with growth form, suggesting that life history can influence molecular evolution at broader genomic scales. Beaulieu et al. (2015) also showed how life history variation in molecular rates can adversely impact divergence-time estimates. Likewise, studies by Bromham and colleagues (2009, 2015) explored how longevity and metabolic rate influence rate variation. Despite these and many other studies (e.g., Beaulieu et al., 2015; Leitch & Leitch, 2012), the specific aspects of life history that impact phylogenomic patterns remain unsettled.

One challenge of studying lifespan as an evolutionary parameter is that lifespan is indirectly connected to several other temporal life history parameters that may impact molecular rates. Maximum lifespan sets a ceiling on average lifespan, maximum generation time, average generation time, time to first reproduction, and average parental age. An increase in the variance of all these life history intervals is also expected as the maximum life span extends. This means that the potential for overlapping generations will tend to increase as life spans increase and generations become shorter. We must also acknowledge the distinction between *maximum potential* lifespan and *average realized* lifespan. This is especially important for trees, for which intrinsic potential lifespans can be extremely long and can differ drastically from realized lifespan, which is largely determined by extrinsic mortality factors. Both lifespan and mortality influence survival and are governed by genetic systems that may be independent or have an underlying connection.

Somatic mutations present an additional challenge for studying the connection between life history and molecular rate, particularly in plants. Trees do not sequester their gametes, which creates the possibility that somatic mutations may accumulate throughout an individual’s lifespan, meaning that longer-lived individuals may carry a diversity of disparate genomes, potentially compensating for lower substitution rates of longer generation times (Burian et al. 2016; Schoen & Schultz 2019). Research on somatic mutations in long-lived organisms is limited, but a few notable exceptions have provided valuable insights. A 234-year-old oak tree (*Quercus robur*) exhibited a surprisingly low number of somatic mutations (Schmid-Siegert et al. 2017). Similarly, an investigation into the massive and ancient Pando aspen clone (*Populus tremuloides*) found modest spatial genetic structure, indicating localized mutation accumulation rather than widespread dispersal over its 16,000- to 18,000-year lifetime (Rozenn et al. 2024). Tissue collected at multiple locations on the corpus of two individuals from different tropical tree species demonstrated that low-frequency somatic mutations are heritable, challenging the assumption that only high-frequency mutations contribute to evolution (Schmitt et al., 2024). Recently, Johannes (2025) suggests that the physical branching structure of trees may limit somatic mutations and their impact on mutation accumulation. While there are still relatively few studies that examine this issue extensively, they collectively highlight that heritable somatic mutations are relatively rare. Still, these mutations can be transmitted to progeny and contribute to population evolution.

Phylogenomic investigations of the past decade have deepened our understanding of long-lived plant evolution. Large-scale sequencing efforts have allowed for the analysis of hundreds, if not thousands, of gene trees and the complexities therein (One Thousand Plant Transcriptomes Initiative, 2019; Smith et al. 2020). One consistent pattern is that long-lived plant species tend to have high rates of gene-tree conflict. Conflict can be acute in long-lived trees; it is generated both by slow substitution rates and short speciation intervals, which create patterns of incomplete allele sorting (i.e., hemiplasy) and by the prevalence of recent and ancient hybridization events (Folk et al. 2018; Pease et al. 2018; Zhang et al. 2021). For example, the Ericales includes lineages that live for more than a thousand years as well as annuals, such as *Primula* species, with their ancestors inferred to be long-lived trees (Carruthers et al. 2024). Gene-tree conflict in Ericales is so extensive that the ancestral relationships remain unresolved (Larson et al. 2020; Carruthers et al. 2024). Fagales includes long-lived species such as *Quercus* and *Fagus* and also exhibits this pattern (Hipp et al., 2020). Given that the external causes of isolation and speciation occur similarly over time (e.g., climatic and geologic events), regardless of the organism’s lifespan, differences in conflict patterns are expected between short- and long-lived species due to the differential rates of substitution.

Here, we address how longevity and time to first reproduction affect population biology and phylogenetics. While traditional population genetics has produced many mathematically sound and powerful tools, there has been relatively less work on the impacts of overlapping generations (although see Felsenstein 1971, Hill 1972, Hill 1979, Nunney 1991, Nunney 1993, Orive 1993, Clark et al., 2024) and long life histories on substitution rates and phylogenetic conflict, and even less when somatic mutations are taken into account. We construct an agent-based simulation to isolate potential causes of lowered substitution rates and increased conflict and then explore the relationship of conflict and effective population size. An agent-based simulation allows us to construct a more realistic model of evolution in which individuals are discrete entities with heritable traits, subject to stochastic birth and death processes, thereby permitting explicit modeling of demographic fluctuations and genetic histories. In this context, we can examine how specific life history properties (maximum age, time to first reproduction, mutation rates, death rate) impact substitution rates and phylogenetic conflict. In addition to this simulation, we examine several empirical cases in more detail to better understand the aspects of their life histories that complicate our ability to reconstruct these phylogenies with confidence.

## Materials and Methods

### Simulations

We constructed an agent-based simulation written in golang and available from http://git.sr.ht/hms/ealdemodor. An initial diploid population is constructed with individuals that consist of one neutral gene with a 1,000 bp sequence. Each diploid individual has two copies of the gene, an age, a maximum age, and a first age at which it reproduces. Each individual is a member of a reproductively isolated population. Individuals within the population share a death rate, a carrying capacity, a branching rate, a meiosis mutation rate, and a somatic/germline mutation rate (referred to as somatic below). The meiosis mutation rate applies at each reproduction event for an individual. The somatic mutation rate is applied each time step.

At the end of each time step, each population is assumed to be at the maximum carrying capacity. At each generation, individuals are removed randomly, following a random uniform distribution, as in |*N*(0.1) · *N*| · *d* where d is the death rate, and *N* is the population size. For each individual, some branching will result in multiple copies of the genes, as in branching in a tree. For each of the genomes within the individual, there is some probability of somatic mutations. If the individual is of reproductive age, it is added to a pool of reproducing individuals. Until the population size reaches the carrying capacity, two random individuals are chosen to reproduce. During reproduction, a random gene is chosen from each parent, mutated based on the mutation rate during meiosis, and a new individual is created and added to the population. Simulations were run for 500,000 generations, with speciation events at 166,666 and 333,333 generations. During speciation, individuals in the speciating population are randomly assigned to one of two new populations, each initially half the size of the original. They will reach carrying capacity in the first generation. When a simulation reached the end of the specified number of generations, the gene sequences were written to a file, after which a phylogeny was created for the living individuals and the starting population (for rooting purposes). With this constructed phylogeny, we calculated the average root-to-tip distance as a measure of substitution rate for the entire simulation. We calculated conflict in two ways. First, we sampled an individual sequence from each population and then constructed a phylogeny (“recalculated”). This was replicated 100 times. Second, we sampled one individual from each lineage from the constructed phylogeny of all individuals, keeping the relationships as they were from the constructed phylogeny (“resampled”). This was replicated 1000 times. In each case, the resulting phylogenies were compared to the known species tree generated by the simulation. We recorded the distribution of ages for each individual at the end of each resulting simulation. We calculated sequence diversity using both *π* (Tajima’s theta) and Watterson’s theta

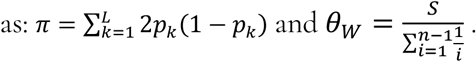

### Simulation runs

Simulations were conducted with the number of generations (500,000), the number of sites (1000), and the birth rate (0.01) remaining constant across runs. Simulations varied or swept the following parameters meiosis mutation rate, somatic mutation rate, branching rate, death rate, maximum age, time to first reproduction, and carrying capacity (detailed in Table 1). In all cases, only one parameter was varied, and all others were kept constant to isolate the effect of the variable under study. Each simulation output consisted of a FASTA-formatted file of terminal gene sequences, along with other output files, to calculate summary statistics. Each parameterization was run for 50 replicates. See Table 1 for more details.

**Table 1.** Simulation parameters and sweep values.

| Parameter Swept | Sweep Values | Fixed Parameters |
| --- | --- | --- |
| Meiosis mutation rate (-m) | 0.0001 - 0.0003 | -l 1e-10, -b 1e-11, -r 10, -x 400 |
| Somatic mutation rate (-l) | 0.0001 - 0.000001 | -m 0.00015, -b 1e-11, -r 10, -x 400 |
| Branching rate (-b) | 0.1 - 0.001 | -m 0.00015, -l 1e-6, -r 10, -x 400 |
| Death rate (-d) | 0.1 - 0.0001 | -m 0.00015, -l 1e-10, -b 1e-11, -r 10, -x 400 |
| Maximum age (-x) | 2 - 1000 | -m 0.00015, -l 1e-10, -b 1e-11, -r 0 |
| Reproduction age (-r) | 0 - 30 | -m 0.00015, -l 1e-10, -b 1e-11, -x 400 |
| Carrying capacity (-n) | 500 - 2500 | -m 0.00015, -l 1e-10, -b 1e-11, -r 10, -x 400 |
| Maximum age (-x) | 2 - 500 | -m 0.0001, -l 1e-10, -b 1e-11, -r 0 |
| Maximum age (-x) | 2 - 500 | -m 0.00012, -l 1e-10, -b 1e-11, -r 0 |
| Maximum age (-x) | 50 - 500 | -m 0.0001, -l 1e-10, -b 1e-11, -r 10 |
| Maximum age (-x) | 50 - 500 | -m 0.00012, -l 1e-10, -b 1e-11, -r 10 |

## Empirical analyses

We examine several empirical datasets to explore the variation in patterns seen in phylogenomic analyses of long-lived species. We examine the Fagales and Cucurbitales, the Ericales, the Caryophyllales, and the Gymnosperms. For each of these datasets, we collected data on longevity, phylogenetic gene tree conflict, and molecular substitutions. The Fagales and Cucurbitales conflict data was obtained from transcriptomic datasets generated as part of Yang et al. (2023). The species-level phylogeny was obtained from Smith and Brown (2018). The Ericales conflict data were obtained from transcriptome datasets as part of Carruthers et al. (2024). The species-level phylogeny also comes from that source and was constructed using Angio353 and Genbank data. The Caryophyllales conflict analyses were performed using the data from Walker et al. (2018) with the species-level phylogeny from Smith et al. (2015). The Gymnosperm datasets, both the transcriptomic dataset for conflict analyses and the species-level phylogeny, were obtained from Stull et al. (2021).

Obtaining maximum longevity estimates pose major challenges for any dataset, but especially for these larger phylogenies. First, there are very few well-characterized datasets of maximum longevity, with exceptions such as Loehle (1988). Larger datasets are available through the Botanical Information and Ecological Network (BIEN; Enquist et al., 2016; Maitner et al., 2018), but their accuracy varies (see Supplementary Material). Furthermore, the general expectation of accuracy is unclear, as longevity can vary significantly depending on environmental factors, genetics, and other variables. Additionally, how exactly those data are obtained, especially for long-lived species, adds more error. With all of this in mind, we attempted to gather rough estimates of longevity for our large phylogenies, understanding that there are likely several sources of error. In each empirical example, we attempted to obtain the available data for each species from the BIEN. In total, there were 4231 records for *maximum whole plant longevity* and *longest whole plant longevity*. For species that were not available but for which the genus was sampled in BIEN, we applied the data to each species in that genus. For taxa for which no information was available, we used doomharvest, applied to each genus with three replicates (https://github.com/FePhyFoFum/doomharvest, Supplementary Material). The results were manually checked for accuracy. Instead of using ages directly, which are prone to being overly precise, we categorized ages into five categories: 0-4, 5-24, 25-99, 100-499, 500-. Estimated values are available here 10.6084/m9.figshare.29087498.

## Results

To examine how lifespan variation shapes evolutionary and population-level dynamics, we simulated allele patterns under a range of demographic and genetic parameters using forward-time, agent-based models (Fig. 1).

**Figure 1.**
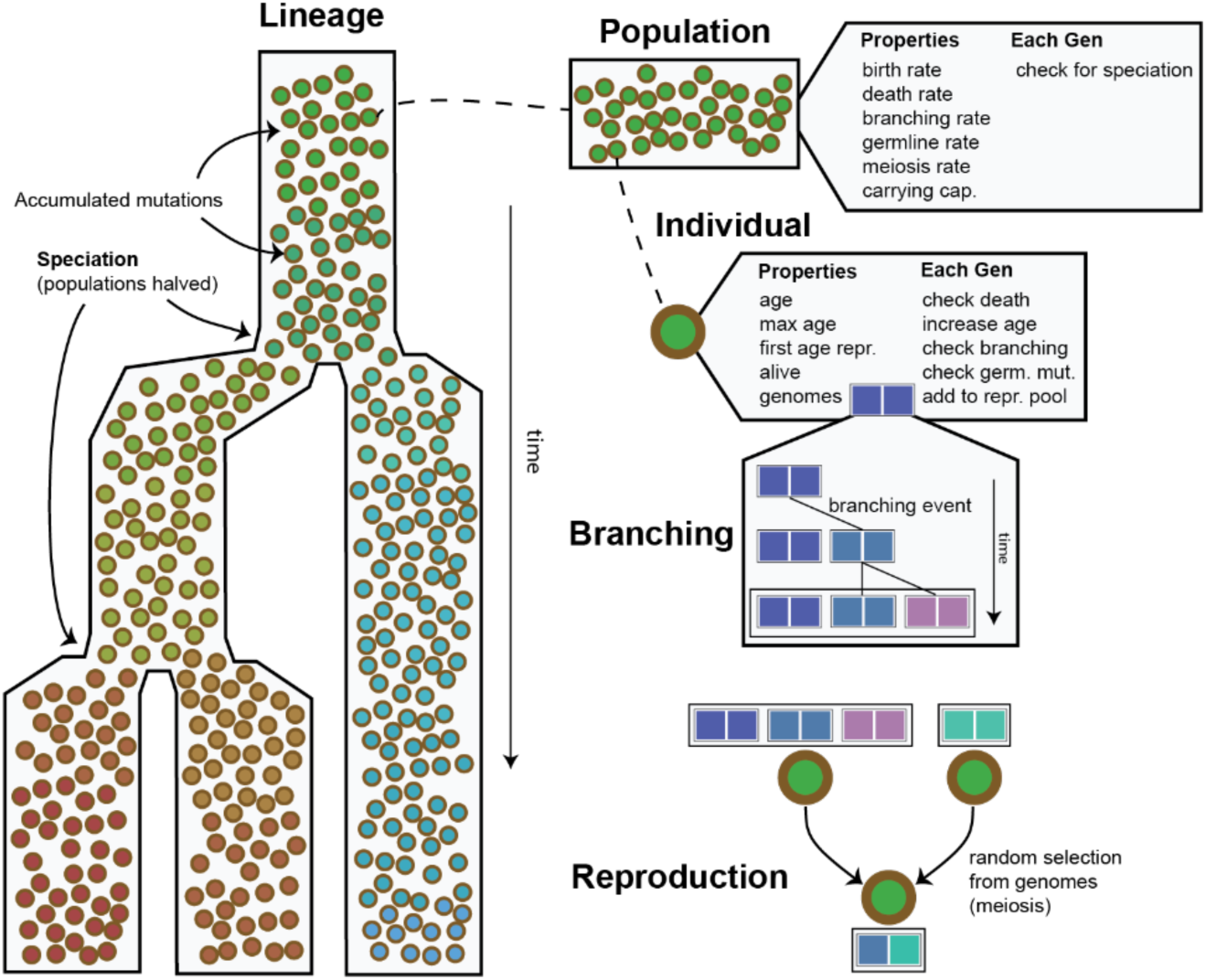
The general design of the agent-based model. A lineage with two speciation events is shown along with populations and their properties, individuals and their properties, a diagram of the branching of genomes and mutations, and reproduction where random haploid genomes are chosen from the branched diploid genomes, meiotic mutations occur and are combined.

We first investigated the effect of maximum individual lifespan on population structure and substitution rates. Populations with shorter lifespans exhibited more rapid generational turnover and faster individual replacement, whereas longer-lived populations accumulated greater age variance and more persistent genome variants. As maximum lifespan increased from 2 to more than 200 years, the average age at death rose, turnover slowed, and substitution rates declined (Figs. 2–3). However, maximum lifespan extensions beyond 500 generations had diminishing effects, suggesting an asymptotic limit beyond which further longevity no longer alters evolutionary dynamics, given the constant death rates under which data were simulated. Notably, longer lifespans also led to increased conflict between inferred allele trees and the known population genealogy (runs 31-34, R^2^=0.91, p=0.045), reflecting the retention of ancestral polymorphism. Interestingly, under high meiotic mutation rates, even long-lived populations showed substitution rates that were nearly saturated, comparable to annuals (Fig. 2; top panels, Fig. 3).

**Figure 2.**
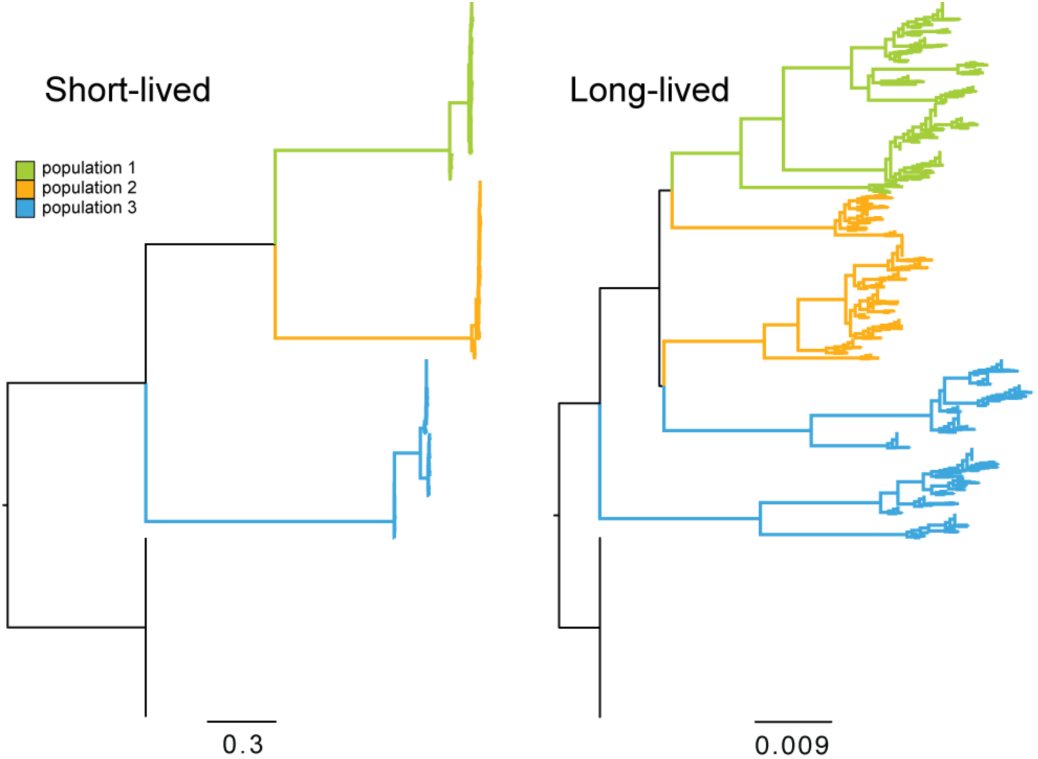
Example phylogenies showing the stark contrast in molecular evolution and tree shape between short-lived and long-lived populations. Individuals within the same population are given the same color. In addition to long-lived populations having lower substitution rates, long-lived populations 2 and 3 are non-monophyletic.

**Figure 3.**
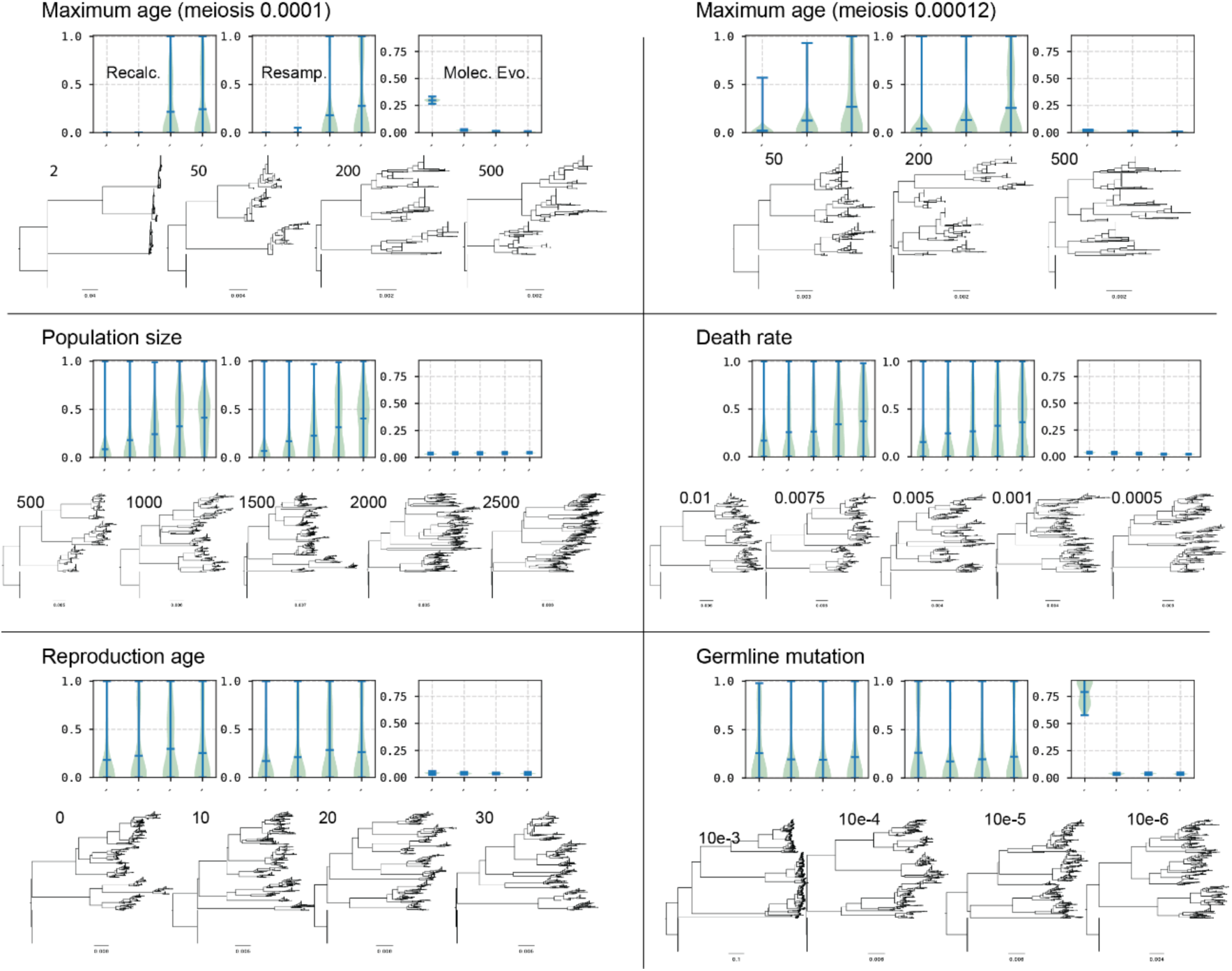
Results from simulation runs. Each set of runs is presented with three different results. The first is the conflict with the species tree when sampling one individual from each population, including the starting population, and recalculating the phylogeny of those four individuals (Recalc.). The second is sampling from the final phylogeny and examining conflict without recalculating the phylogeny (Resamp.). The third is calculating the average tip-to-root height to measure molecular evolution. The numbers are referenced in Table 1 for additional information on parameters. The trees below and the associated number correspond to each entry in the graph. Additional results are presented in the supplement.

Age at first reproduction also influenced genome evolution. Earlier maturation (e.g., at age 0) slightly increased mutation accumulation and substitution rates (from 0.04 with maturation of 0 to 0.035 with maturation at 30), while delayed maturation (e.g., age ≥10) slowed population turnover. Gene-tree conflict was reduced under earlier maturation, from a mean of the proportion of conflicting trees of 0.18 with age 0 at maturation to 0.25 with age at maturation of 30. However, these effects were consistently weak relative to lifespan changes, where conflict went from 0 to 0.24 with maximum age from 0 to 500 (all with maturation of 10 years).

Variation in the per-generation death rate altered the population’s turnover dynamics. Higher death rates (≥0.01) increased turnover, reduced average lifespan, and produced younger-skewed age distributions (see Supplement). These conditions also elevated substitution rates (from 0.023 with death rates of 0.0005 and 0.038 with death rates of 0.01) and reduced gene-tree conflict (halving conflict from 0.36 with death rates of 0.0005 to 0.15 with death rates of 0.01). As death rates decreased, substitution rates plateaued, reaching asymptotes at 0.0001 deaths-per-generation-per-capita.

Simulations across a range of carrying capacities revealed two distinct and general regimes. Smaller populations (e.g., *n* ≤ 500) exhibited less gene-tree conflict (0.06 proportion of conflicting trees) due to faster lineage turnover, regardless of time interval. In contrast, larger populations (*n* ≥ 1500) maintained higher levels of standing variation and ancestral polymorphisms, and gene-tree conflict (0.22 proportion of conflict for *n* = 1500 and 0.41 proportion of conflict for *n* = 2500; runs 26-30, R^2^=0.98, p=0.001). Substitution rates were not dramatically influenced (from 0.03-0.04 with *n* = 500 and 2500, respectively), even with extremely small population sizes, when the maximum age was still high (e.g., 200-500).

Finally, somatic mutation rates had a strong influence on genome divergence. Elevated somatic mutation rates, and especially increased germline mutation rates, greatly amplified substitution rates and sequence diversity. When germline rates exceeded approximately 0.0001 mutations per generation per individual, they could compensate for the otherwise slow rates observed in long-lived populations. However, such high rates likely exceed those observed in most empirical systems. It is also important to consider that, as with real systems, the simulations have a “spatial” component to somatic mutations due to the addition of branching. This could also impact the results and should be explored further.

### Empirical results

We examined life history, molecular evolution, and conflict in four major clades of seed plants: Caryophyllales, Ericales, Fagales, and Cucurbitales, as well as Gymnosperms. The Caryophyllales have life spans ranging from the long-lived cacti to annuals, such as those in the Nyctaginaceae (Smith et al., 2021). The most conflict in the transcriptomic data is within the longer-lived cacti lineages and along the backbone, which is reconstructed as somewhat intermediate in lifespan (Yang et al. 2021). It is essential to note that while longevity has been correlated with shifts in molecular rates, rate changes have also been linked to transitions into different climates within the Caryophyllales (Smith et al. 2021). Here, a higher proportion of conflict correlates with longevity (Spearman ρ=0.356, p<0.0001) as well as with lower log-substitution rates (Spearman ρ=−0.104, p<0.0001, Fig. 4). The Spearman ρ for branch lengths are low, but that is to be expected with these large complex clades where certainly many processes are impacting both conflict and rates of molecular evolution; we are likely finding only one of many correlates.

**Figure 4.**
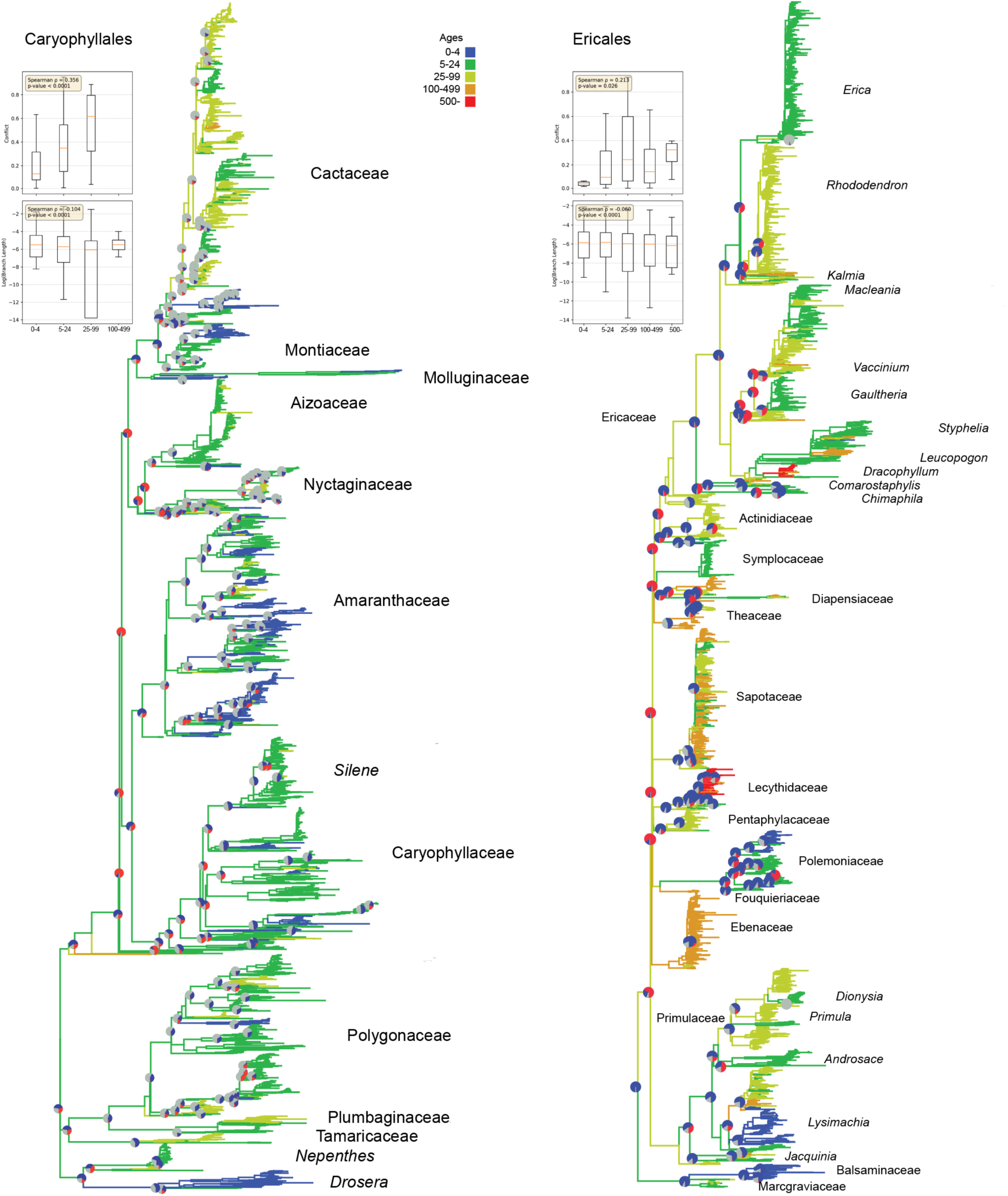
Conflict and molecular evolution in the Caryophyllales (left) and Ericales (right). Pie charts show conflict (red), concordance (blue), and lack of information due to low bootstraps (gray).

Ericales includes shorter-lived species such as *Impatiens* and several long-lived species, such as *Pouteria* (Sapotaceae), which can live for several hundred years. The Ericales also include many long-lived tree genera, such as those in the Lecythidaceae and *Dracophyllum* (Ericaceae). The backbone of the Ericales has extensive gene-tree conflict and tends to be reconstructed as a long-lived tree. We find a correlation between conflict and longevity (Spearman ρ=0.213, p=0.026); short branch lengths (log-substitution rates) also correlate with longevity (Spearman ρ=−0.060, p<0.0001; Fig. 4). As with the Caryophyllales, Spearman ρ for branch lengths are low. The results are similar to those found by Carruthers et al. (2024), who examined life history using a broader categorization of habit.

The Fagales and Cucurbitales demonstrate a major contrast in life-history and longevity (Fig. 5). The Fagales contain some of the longest-lived angiosperm tree species within the flowering plants, and the Cucurbitales include many short-lived species and annuals. There was a stark contrast in molecular substitution rates between these two orders. While we have fewer Cucurbitales nodes with conflict information than Fagales, the average conflict is much higher in Fagales (0.423) than in Cucurbitales (0.088). We find a strong correlation between conflict and longevity (Spearman ρ = 0.387, p < 0.0001); short branch lengths (log substitution rates) also correlate with longevity (Spearman ρ = −0.352, p < 0.0001). While the results are strong within this clade, notably, there are fewer nodes with conflict information available for the herbaceous Curcurbitales.

**Figure 5.**
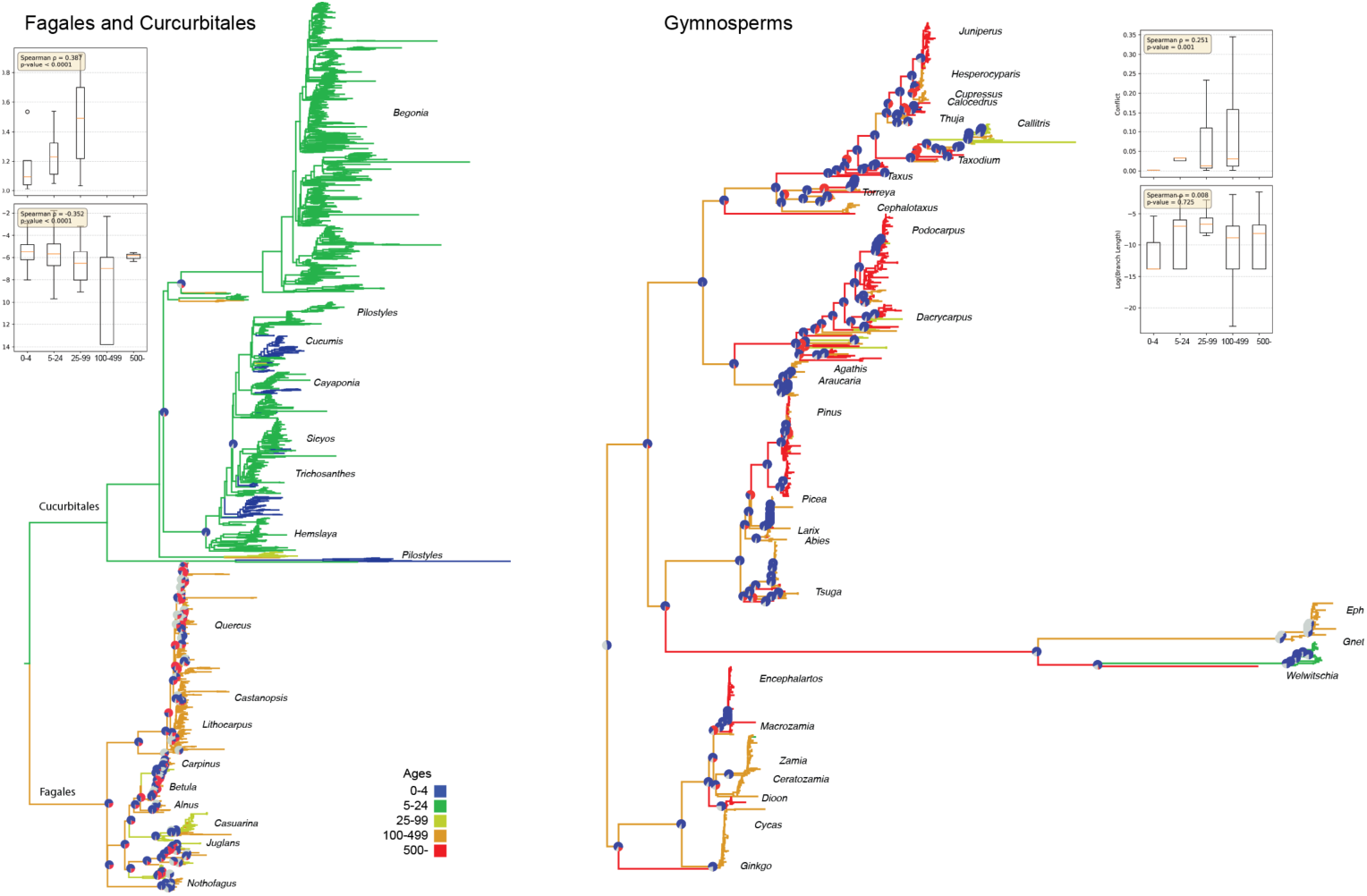
Additional empirical examples of conflict and molecular evolution in the Fagales + Curcurbitales and Gymnosperms. Pie charts show conflict (red), concordance (blue), and lack of information due to low bootsraps (gray).

Gymnosperms include some of the longest-lived plant species on earth. *Pinus longaeva* (Pinales) can live several thousand years, and *Welwitschia mirabilis* (Gnetales) can live more than 1500 years (Fig. 5). In contrast, other Gnetales are relatively short-lived species and their branch lengths are much longer than the remainder of the Gymnosperms. Gene-tree conflicts vary significantly across the clade, with several long-lived species exhibiting high conflict within the Cupressaceae, Araucariales, and Pinales (average conflict 17% and maximum conflict of 49%). In contrast, the Gnetales have very little conflict, with only one conflicting gene tree. As with the other datasets, we find a correlation between conflict and longevity (Spearman ρ = 0.251, p = 0.001). However, short branch lengths (log substitution rates) did not correlate with longevity (Spearman ρ = −0.008, p = 0.725). This test is likely insignificant because there are few nodes with young species (only three taxa with a longevity of less than 55 years).

## Discussion

Our results demonstrate that fundamental aspects of life history can greatly impact phylogenetic conflict patterns and influence expectations of population genetic evolution. Our primary finding, supported by both simulations and empirical data analysis, is that long-lived species more often exhibit higher conflict and fewer molecular substitutions than short-lived species. The impact of lifespan variation on conflict and molecular substitution rates depends on several parameters. Extreme longevity results in low substitution and deep divergences within the population, mimicking larger population sizes with higher incomplete lineage sorting (Fig. 2). The primary cause is overlapping generations, where reproductively active long-lived individuals cause “old” alleles to persist in the population both by their presence in the population and by contributing them to new offspring. If a potentially long-lived population has higher realized death rates, then its allele and conflict patterns will mimic those of a shorter-lived population, depending on the death rate and age at maturity. When the age at maturity is older, substitution rates tend to be lower, and conflicts tend to be higher. Somatic mutations and branching have the effect of increasing the apparent substitution rate when they are high, though there is still little evidence for high rates of somatic mutations in extant populations (citations). Population size differences have the expected effect of larger populations of long-lived species having more conflict.

These findings have several implications for the long-standing discussions about molecular rate heterogeneity in angiosperms (Beaulieu et al., 2015; Smith & Beaulieu, 2024). Those conversations have proposed several potential biological mechanisms that might influence generation time and effective population size to explain the differences in molecular evolutionary rates between short- and long-lived species (e.g., DNA repair, metabolic rate, and life history; Martin and Palumbi 1993, Bromham 2009). In plants, life history axes such as annual versus perennial or herbaceous versus woody have been thought to be particularly important in determining evolutionary rates (e.g., Gaut et al. 1997, 2011, Andreasen and Baldwin 2001, Smith and Donoghue 2008, Lanfear et al. 2013). While rates of molecular evolution have been suggested to be strongly correlated with life history, the mechanisms have not been adequately examined (Smith and Donoghue 2008, Yang et al. 2012). The simulations presented here demonstrate how overlapping generations, age-structured reproduction, and retention of ancestral polymorphisms can reduce the rate at which mutations become fixed, thereby lowering substitution rates. Furthermore, the gene-tree conflict seen in some long-lived lineages may be explained by the interaction between speciation time, longevity, and population size. This is consistent with observed gene-tree discordance in groups such as Fagales (Hipp et al., 2020) and Ericales (Larson et al., 2020; Carruthers et al., 2024), where long-lived lineages exhibit extensive phylogenetic conflict.

While these simulations demonstrate that several life history and demography parameters can impact phylogenetic patterns, these parameters can exhibit strong variation within lineages and even among populations of the same species. This expected variation is clear from the empirical examples, where short branch lengths can exhibit relatively higher conflict (e.g., Ericales and Fagales) or relatively lower conflict (e.g., Gymnosperms and Caryophyllales; Figure 4-5). The variation at these large scales likely reflects the extensive variation in life-history and other properties among lineages. Variation within lineages may be harder to detect at phylogenetic scales. Nevertheless, our simulations suggest patterns we might examine with the right datasets.

Of the several parameters we examined, death rate emerges as one of the more straightforward in interpreting its effects. In empirical systems, populations of long-lived species are likely to vary significantly in death rate due to differences in local disturbance or ecological relationships. We demonstrate in simulations that populations with higher death rates are likely to exhibit patterns more similar to those of shorter-lived species, with implications for substitution rates, population dynamics, and phylogenetic conflict. In empirical systems, the spatial variation in death rate will complicate these patterns. The impact of integrating populations with spatial variation in death rates on the phylogenetic patterns observed here is not well understood, although these issues have been discussed in relation to age distributions in ecological studies (Wright et al. 2003; Petit and Hampe 2006). Spatial variation in death rate and its effects should be examined further in both empirical and simulation studies.

Another consideration for further examination is the time between speciation events. Here, we demonstrate the impacts of life history on substitution rates and therefore branch lengths. However, the time between speciation events will be another significant factor in the realized branch lengths. Along with allele diversity, speciation interval is one of the two classic parameters of incomplete allele sorting (Hudson 1983, Tajima 1983), where a longer time between speciation events leads to a lower probability of sampling allele conflict in descendant species. The speciation rate is a population- or metapopulation-level process whose intrinsic and extrinsic causes are generally stochastic and decoupled from the lifespan of organisms. For extraordinarily long-lived species, one might imagine that a particularly short speciation interval might be on the order of only a handful of maximum lifespan generations. In contrast, a short-lived species with a long inter-speciation interval might experience millions of generations between splits. Under genetic reproductive isolation, we expect the molecular divergence to be proportional to the speciation rate. However, the stochastic nature of genetic barriers means that the variance provides only a loose ceiling on this pattern. Therefore, the relative timing of generations, lifespan, and speciation events may exert a strong effect on phylogenetic patterns, as shown here.

While we have addressed some of the standing questions facing life history and molecular evolution, many more remain unanswered. For example, the simulations only include two evenly spaced speciation events to simplify the model, resulting in three final reproductively isolated populations. The frequency and rate of speciation both have a significant impact on phylogenetic patterns and could be explored further in conjunction with the parameters examined in this study. We also only consider populations that share life history traits. This was motivated to isolate the impacts of individual parameters on molecular variation. However, life history shifts occur across the plant tree of life at varying frequencies. The effects of these changes on the patterns of rate variation and gene tree conflict require additional examination. Additionally, we only look at a one-thousand-base-pair gene region. To better reflect our data collection efforts, expanding the number of gene regions to hundreds or thousands would be enlightening. Each extension can be done within the current framework discussed in the methods and implemented in our code. Other future axes of complexity beyond the simulation framework would also be fascinating to explore, such as spatially explicit patterns of molecular variation.

This study presents an agent-based model to explore the impacts of life history and population dynamics on molecular evolution. Mechanisms exist to examine certain components and variables within a more traditional mathematical framework of population genetics. However, those traditional frameworks are limited in fully accommodating the complexity of some of the elements explored here. For example, while the literature on life history and molecular evolution frequently discusses longevity, a potential outcome of longevity—overlapping generations—is rarely considered. Additionally, the potential for somatic mutations that may accumulate over a long-lived plant’s lifetime to contribute to a population’s molecular evolution adds a significant layer of complexity. Agent-based models allow for the incorporation of unconstrained development of models with realistic parameterizations. Of course, this, in turn, comes with its challenges, as the added complexity may overwhelm the focus of the question at hand. Striking a balance between adding variables that are essential to addressing the hypotheses in question and adding superfluous variables can be a difficult task.

While many extensions to the modeling framework await exploration, our findings also highlight facets of available data that are underexplored. Genomics datasets range from those with species-poor coverage and broad sampling to those with extensive resampling of a single species. Several projects have aimed to greatly expand the sampling of genomic datasets, and these resources will undoubtedly become available. However, in the meantime, there are several barriers to using these data for addressing some of the questions explored here. Datasets focusing on population genetics are invaluable for exploring phylogeographic patterns and recent population dynamics. However, these datasets are often too limited in geographic or phylogenetic scope to capture transitions in life history or other complexities. In contrast, systematic datasets rarely sample more than one individual for each species. An exciting avenue lies in waiting for datasets that span these typical sampling boundaries.

While exploring and extending these simulations will provide additional insights into the impacts of life history on broad plant evolution, more in-depth empirical analyses would also be essential. Unfortunately, and as discussed in the methods and supplement, data on maximum longevity are particularly scarce for both genera and species. Efforts to improve these datasets, along with the inherent uncertainties in the data, would be highly fruitful for furthering our understanding of the evolution of life histories.

## Acknowledgments

SAS and IBH were supported by NSF-DEB IntBio #2217116. JBP was supported by NSF-DEB IntBio #2217117. JMB was supported by NSF-IOS #2409451. Research reported in this publication was supported by the National Institute of General Medical Sciences of the National Institutes of Health under Award Number R35GM137919 (awarded to G.S.B.). The content is solely the responsibility of the authors and does not necessarily represent the official views of the NIH. SAS would like to thank the Smith lab for comments and discussions.

## Author contributions

SAS wrote the code and conducted analyses. SAS, JBP, and JMB initially conceived the study. JBP and GSB assisted with population and simulation analyses. TSY, YYY, GWS, TC helped analyze the empirical datasets. IBH and WNW assisted with data collection and doomharvest processing. All authors contributed to the manuscript.

## Data availability

The data that supports the findings of this study are available at 10.6084/m9.figshare.29087498. Other data have been previously published and cited in the text.

