## Supplemental Materials for "Longevity in plants impacts phylogenetic and population dynamics"

**Supplementary Figures**

Age distributions under different scenarios.

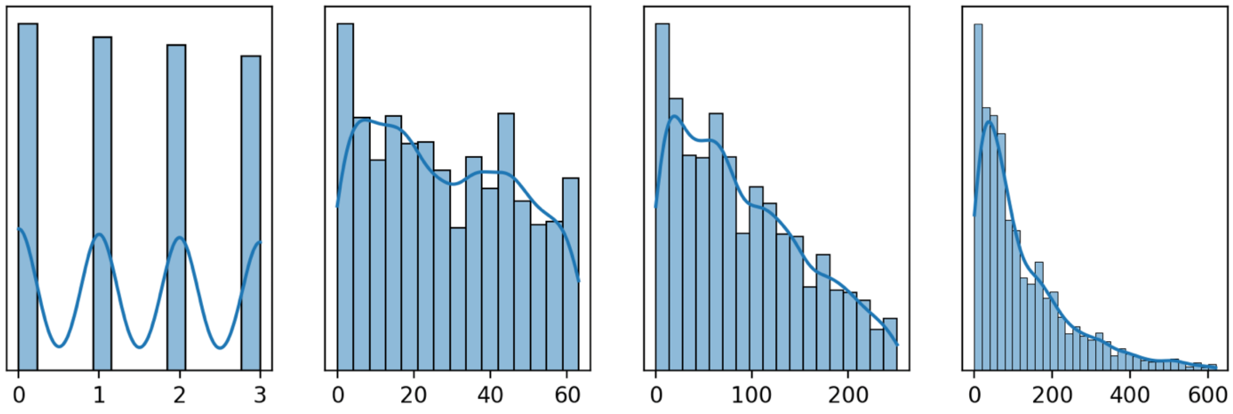


Supplementary Figure 1. Age of individuals from populations at the end of the simulation with maximum age varying from 2, 50, 200, and 500 from left to right.

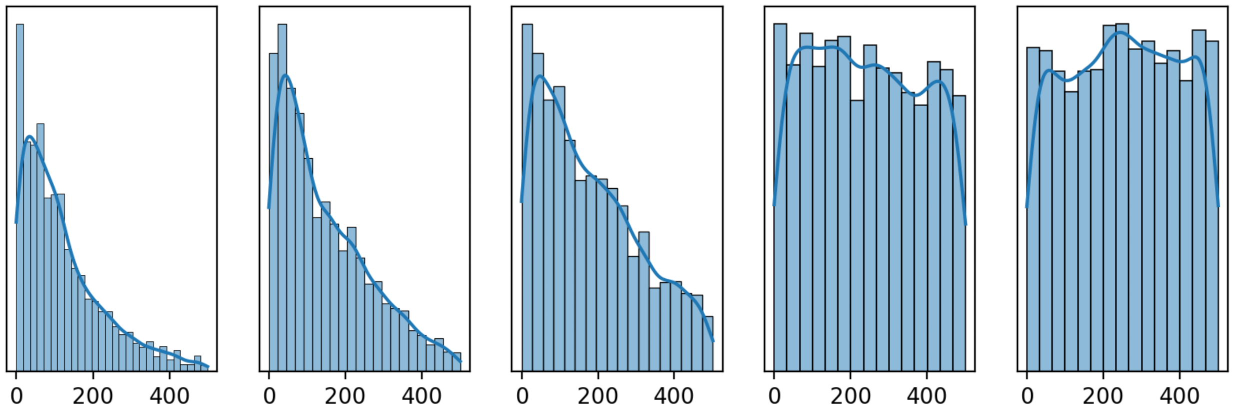


Supplementary Figure 2. Age of individuals from populations at the end of the simulation with death rates varying from 0.01, 0.0075, 0.005, 0.001, and 0.0005 from left to right.

**Supplementary Methods**

**Aggregating Life History Data with doomharvest**

Doomharvest is a Python package that queries GPT-4o (ChatGPT) for natural history data. The package uses an initial prompt template to query GPT-4o multiple times (a default of three times) to find the character state of a trait for a given taxon, uses a second prompt as a quality check, and then exports a filtered summary of the results.

The initial prompt assigns an "expert" role (to encourage use of technical sources), provides context (whether the information queried relates to reproductive or vegetative morphology), provides format instructions (an easy to compile, numerical JSON dictionary format), requires reputable sources, and asks for a self-assessed confidence value for the results. The confidence values can be between 1-10 (where 10 represents the maximum confidence in results) or -1 for instances where GPT-4o cannot provide an answer. When aggregating information on discrete characters, this initial prompt also provides the bins for the character state options as well as potential synonyms, vocabulary, or phrases that refer to each character state. Results from the initial prompt that have a low confidence score (1-5 or -1) are coded as unknown ("?") in the final outputs.

The second prompt asks if a reliable source of information on the trait for the given taxon exists. The output is either 1 (reliable information on the trait exists for this taxon) or -1 (reliable information on the trait does not exist for this taxon). Like the initial prompt, the sanity check asks GPT-4o and the user to play expert roles. To account for biases related to taxonomic rank, the doomharvest package checks if information on a trait is reported at family, genus, and species level; if a trait is not reported at a higher taxonomic rank, it is treated as unknown ("?") for subsequent/lower taxonomic ranks. After filtering out low confidence results from the first prompt and -1 results from the second prompt, the results of prompt replicates are averaged.

**Prompt Templates**

For discrete traits, the initial prompt template is as follows:

*I am aggregating data for a research project that is looking at the biology of angiosperm plant species. You are a specialist in botanical taxonomy. In regard to CONTEXT, I need you to tell me whether the taxon that the user provides has INSERTTEXT1 in a JSON dictionary like this: {{'taxon_name': {{'value', 'confidence': confidence_in_answer, 'sources': ['source_1', 'source_2', '...'],'annotations': 'annotation'}}}}. Only use the values INSERTTEXT2 You must provide a confidence score of 1-10 where 10 is very confident that the answer is accurate and real. Return only the JSON dictionary with no explanation. If you don't know the answer, set the confidence to -1. Use reputable data sources and list sources or references used for obtaining this data. Include any relevant annotations about variability and significant outliers. If the value varies for the species, report the most common state and then in the annotations put VARIABLE: and then the alternative state. The taxon is NAMEOFTAXON.*

The bracketed text varies depending on the trait being queried. "CONTEXT" can be adjusted depending on the category of trait (e.g., "floral morphology", "vegetative morphology", "fruit and seed morphology"). "INSERTTEXT1" describes the trait being queried, and "INSERTTEXT2" describes the bins for the character states as well as phrases and synonyms for traits, if applicable. The taxon provided ("NAMEOFTAXON") can be for any clade or taxonomic rank. For continuous traits, the initial prompt template is identical to that used for discrete traits, but the phrase, "Only use the values" is excluded.

The second prompt template is as follows:

*I am aggregating data for a research project that is looking at the biology of angiosperm plant species. You are a specialist in botanical taxonomy. Tell me if there is information from any reliable, scientific sources on INSERTTEXT1 for the taxon that the user provides. Format an answer in a JSON dictionary like this: {{'taxon_name': {{'value', 'sources': ['source_1', 'source_2', '...'],'annotations': 'annotation'}}}}. Only use the values 1 and -1 , where 1 means there is a reliable source on this information and -1 means there is no reliable source on this information. Only report a source if you can find a reliable source. The taxon is NAMEOFTAXON.*

**Factors impacting accuracy**

***Trait name***

Results are less accurate for traits that use the same descriptive terms (e.g., "sessile anthers" and "sessile leaves" both use the word "sessile"), which includes traits related to phyllotaxy (e.g., "spirally arranged leaves" vs. "spirally arranged petals"), fusion (e.g., "connate petals" vs. "connate stamens"), or convergent morphologies (e.g., "petaloid staminodes" vs. "petaloid styles"). This semantic confusion can be combatted by including additional context in prompts, providing which terms are synonymous, which are inequivalent, and example phrases of the term's usage. For example, *Iris*, which lacks staminodes, was often reported by GPT-4o as having petaloid staminodes; *Iris* does not have petaloid staminodes but has petaloid styles. Including the statement, "petaloid staminodes are not the same as petaloid styles," improved results.

Some traits are given different names depending on the taxonomic group they are used in (e.g., "costa" in palms and ferns vs. "midvein" in most angiosperms) or whether they are included in a compound structure (e.g., "floret" vs. "flower"). Providing synonyms and example phrases can mitigate errors from this.

***Taxonomic level and accessibility of information***

Results at the genus and family level were more accurate than results at the species level. Since many plant species share the same epithet, a target species may be conflated with another species from an unrelated clade.

Although GPT-4o is trained on massive datasets, it is not guaranteed that these datasets contain information on a given taxon. Plant families are more easily found in textbooks, floras, and databases compared to their constituent genera. Similarly, genera are more likely to have widely accessible information reported on them than their constituent species. Even if detailed descriptions that thoroughly describe the features being queried exist for a given species, these descriptions may be in printed monographs or paid online sources, which models like GPT-4o would not be trained on. To mitigate this, we adapted the doomharvest package to assume that a species has no information on a trait (treated as "?") if it is not reported for its genus. Similarly, the character state for a genus is treated as "?" if information cannot be found about this character for the family level. For similar accessibility reasons, results for commercially important taxa and common, temperate taxa were better than results for taxa from regions of the world with historically less documentation, such as the Neotropics.

***Example sources***

The initial prompt calls for reputable sources but does not call for specific sources. We experimented with providing examples of specific sources such as Flora of North America (www.floranorthamerica.org), DELTA (www.delta-intkey.com), and World Flora Online (www.worldfloraonline.org), but these queries often produced false positives. For a few traits, (e.g., unisexual vs. bisexual flowers) providing sources (i.e., including the phrase "The best sources are these websites:" and providing links) improved accuracy. This is only recommended for difficult traits.

**Recommendations for users**

When picking a trait to harvest, consider its synonyms and if other traits use similar verbiage. Provide information about synonymous terms or phrases that do not reflect the targeted trait. If one query is not successful, try breaking the query down into multiple queries or rephrasing the query as a simpler, binary question (e.g., instead of "Does the taxon have X or Y?" ask, "Does the taxon have X?"). Confidence values may be overinflated but using them as cutoffs can help refine results. Test queries at multiple taxonomic levels to check for accuracy; the same biases affecting data repositories may affect results from GPT-4o. Results for taxa that are more commonly reported in online sources—including higher taxonomic levels (genera and family), agriculturally important, common, or ornamental taxa—will likely be better than others. Averaging results from multiple runs of a query can improve results. Finally, we recommend iterative refinement. Prompts will likely require repeated tests and multiple stages of revision before they can be useful for aggregating information. Manually checking results or comparing results to a ground truth dataset is important to help find issues with prompts before harvesting broadly.

**Accuracy of "maximum age" for plants**

As discussed in the main text, there are few major sources for the maximum age of plants. One such source is the study by Loehle (1988). We compared the values of maximum mortality from this study to the values from BIEN traits "maximum longevity" and "longest longevity" as well as to the results from doomharvest.

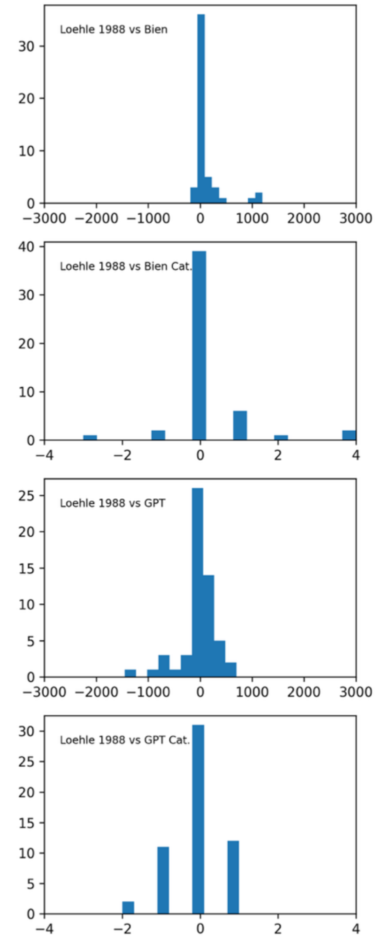


Supplementary Figure 3. Results comparing both the age estimate and the category estimate between Loehle (1988) data and BIEN and doomharvest for genera. Positive values indicate higher values for Loehle (1988).

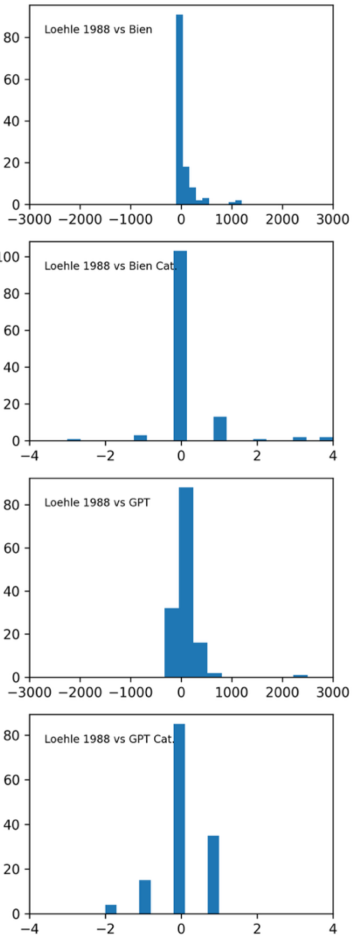

Supplementary Figure 4. Results comparing both the age estimate and the category estimate between Loehle (1988) data and BIEN and doomharvest for species. Positive values indicate higher values for Loehle (1988).
